# Nutritionally responsive PMv DAT neurons are dynamically regulated during pubertal transition

**DOI:** 10.1101/2025.02.03.636271

**Authors:** Cristina Sáenz de Miera, Nicole Bellefontaine, Marina A Silveira, Chelsea N Fortin, Thais T Zampieri, Jose Donato, Kevin W Williams, Cristiano Mendes-da-Silva, Laura Heikkinen, Christian Broberger, Renata Frazao, Carol F Elias

**Author notes:** Corresponding Author: Carol F. Elias, Ph.D., North Campus Research Complex B25-3682, 2800 Plymouth Road, Ann Arbor, MI 48109.

## Abstract

Pubertal development is tightly regulated by energy balance. The crosstalk between metabolism and reproduction is orchestrated by complex neural networks and leptin action in the hypothalamus plays a critical role. The ventral premammillary nucleus (PMv) leptin receptor (LepRb) neurons act as an essential relay for leptin action on reproduction. Here, we show that mouse PMv cells expressing the dopamine transporter (DAT) gene, *Slc6a3* (PMv^DAT^) form a novel subpopulation of LepRb neurons. Virtually all PMv^DAT^ neurons expressed *Lepr* mRNA and responded to acute leptin treatment. Electrophysiological recordings from DAT^CRE^;tdTomato mice showed that PMv^DAT^ cells in prepubertal females have a hyperpolarized resting membrane potential compared to diestrous females. *Slc6a3* mRNA expression in the PMv was higher in prepubertal than in adult females. In prepubertal females *Slc6a3* mRNA expression was higher in overnourished females from small size litters than in controls. Prepubertal *Lep^ob^* females showed decreased PMv *Slc6a3* mRNA expression, that recovered to control levels after 3 days of leptin injections. Using a tracer adenoassociated virus in the PMv of adult DAT^Cre^;Kiss1^hrGFP^ females, we observed PMv^DAT^ projections in the anteroventral periventricular and periventricular nucleus (AVPV/PeN), surrounding Kiss1^hrGFP^ neurons, a population critical for sexual maturation and positive estrogen feedback in females. The DAT^CRE^;tdTomato projections to the AVPV were denser in adult than in prepubertal females. In adults, they surrounded tyrosine hydroxylase neurons. Overall, these findings suggest that the DAT expressing PMv^LepRb^ subpopulation play a role in leptin regulation of sexual maturation via actions on AVPV kisspeptin/tyrosine hydroxylase neurons.

**Significance Statement:** Women with excess or low energy stores (*e.g.,* obesity or anorexia) have reproductive deficits, including altered puberty onset, disruption of reproductive cycles and decreased fertility. If able to conceive, they show higher risks of miscarriages and preterm birth. The hypothalamic circuitry controlling the interplay between metabolism and reproduction is poorly defined. Neurons in the ventral premammillary nucleus express the leptin receptor and play a key role in the metabolic control of reproduction. Those neurons are functionally and phenotypically heterogeneous. Here we show that a subset of leptin-sensitive neurons co-expresses the dopamine transporter (DAT), is dynamically regulated during pubertal transition and with nutrition and projects to brain sites relevant for sexual maturation.

**Graphical abstract:** 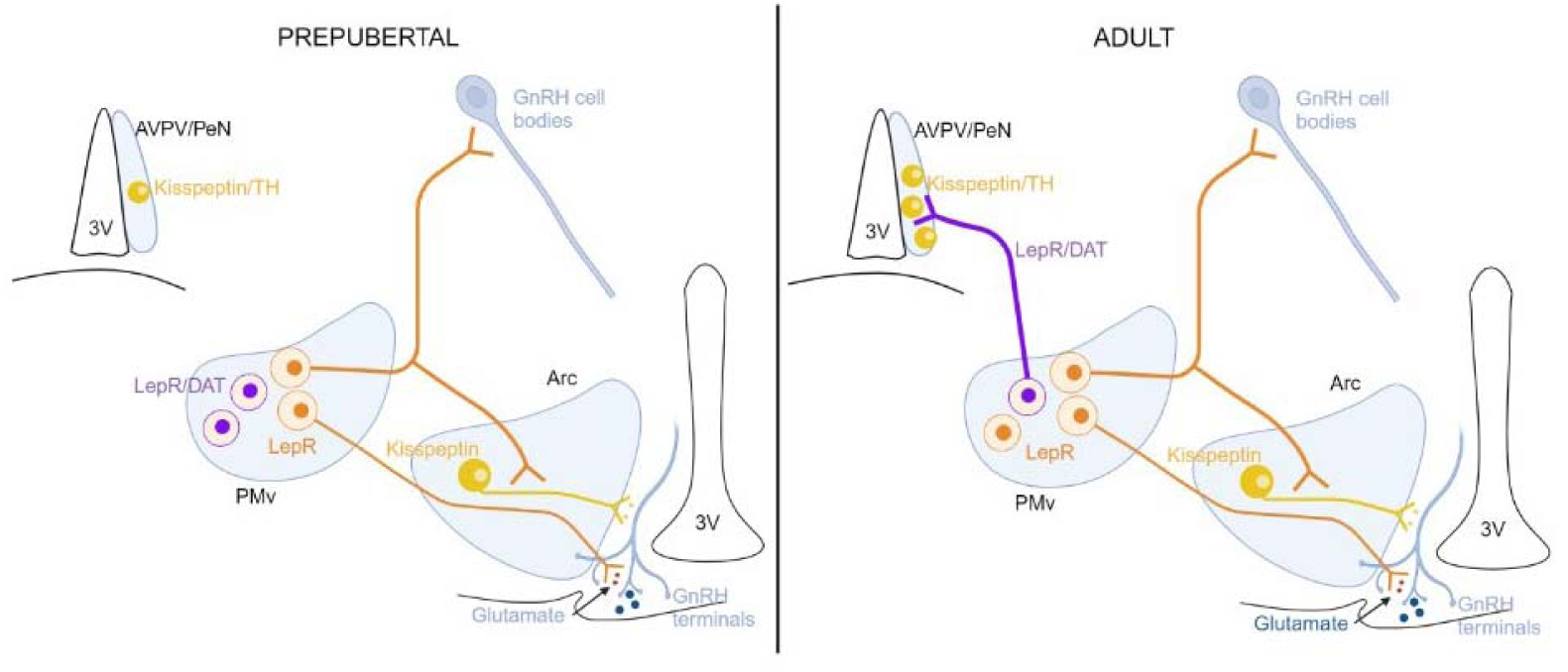

The ventral premammillary nucleus of the hypothalamus plays an essential role in the metabolic control of reproduction. Puberty brings large changes to a subpopulation of PMv^LepRb^ cells expressing the dopamine transporter (PMv^DAT^). DAT gene expression is higher in prepubertal than in adults and is regulated by leptin in prepubertal females. Dynamic projections from PMv^DAT^ cells contact the kisspeptin and tyrosine hydroxylase (TH) populations in the AVPV/PeN during puberty, a critical time for the appearance of these cells in the AVPV/PeN.

## Introduction

Pubertal development and the maintenance of reproductive function are disrupted in states of negative energy balance or excess energy reserve (1–3). If energy stores are low, puberty is delayed, the reproductive cycles are prolonged, and sub- or infertility ensues (2–4). High adiposity, on the other end, induces earlier pubertal development and decreased fertility in adult life (5–7). The cross-talk between metabolic and reproductive functions is orchestrated by a complex neuronal network modulated by circulating hormones and metabolic cues (8, 9). Among them, leptin has critical roles (10–12). Leptin signaling-deficient subjects develop obesity and remain in an infertile prepubertal state (13–15). In mice, direct leptin actions only in the brain are sufficient to normalize body weight, induce puberty and maintain fertility (10, 14, 16).

The ventral premammillary nucleus (PMv) contains a dense collection of leptin receptor (LepRb) neurons, and is recognized as an important hypothalamic site in the metabolic control of reproductive function (4, 16–21). Bilateral lesions of the PMv disrupt estrous cycles and the ability of leptin to increase luteinizing hormone secretion after fasting (4). Endogenous restoration of LepRb exclusively in PMv neurons rescues pubertal maturation and fertility in LepRb *null* female mice (16), while activation of PMv LepRb neurons is sufficient to induce LH release even in normally fed female mice (20). The PMv LepRb neurons, however, do not comprise a homogeneous population, *i.e.,* about 75% depolarize and 25% hyperpolarize in response to leptin (22), but their seemingly dissociated nature and function are poorly understood.

The PMv neurons are mostly glutamatergic and innervate brain sites associated with reproductive control, sending direct inputs to kisspeptin and gonadotropin-releasing hormone (GnRH) neurons (16, 18, 19, 21, 23). A subset of PMv neurons also expresses the dopamine transporter (DAT), a membrane protein associated with dopamine reuptake at presynaptic terminals (24–26). PMv^DAT^ neurons are unique in the sense that they show seemingly undetectable levels of tyrosine hydroxylase (TH) and dopamine release to specific brain sites (25, 27). Manipulation of PMv^DAT^ neuronal activity has shown an action in male social behavior, inter-male and maternal aggression and maternal behaviors (25, 26, 28, 29). However, the role of PMv^DAT^ neurons in female reproductive physiology has not been described, and whether they participate in the metabolic control of reproductive function is unknown.

In this study, we show that DAT is expressed in a subpopulation of PMv^LepRb^ neurons. *Slc6a3* (*DAT*) mRNA expression is higher in prepubertal than in adult females and it is increased in overnourished prepubertal females. We also show that PMv^DAT^ neurons project to and make apparent contacts with kisspeptin neurons in the the anteroventral periventricular nucleus (AVPV) in adults, but not in prepubertal mice.

## Methods

### Experimental animals

All procedures were carried out in accordance with the National Research Council Guide for the Care and Use of Laboratory Animals, and protocols were approved by the University of Michigan IACUC (PRO000010420); and in accordance with the European Community Council directive of November 24, 1986 (86/609/EEC) and had received approval by the local ethical board, *Stockholms Djurförsöksetiska Nämnd.* Mice were held under a 12h:12h light:dark cycle (lights on at 6 am), temperature-controlled at 21-23 °C, and fed *ad libitum* on a low-phytoestrogen diet (Envigo 2016 diet) and a higher protein and fat phytoestrogen reduced diet 2019 (Envigo 2019 Teklad diet) when breeding. Strains of mice used were a line expressing Cre-recombinase under the *Slc6a3* promoter (DAT^Cre^, JAX®; Stock 006660) (30), and (31) only for female electrophysiology, a ROSA26 stop-floxed tdTomato reporter mouse line (tdTomato, JAX®; Stock 007914), mice expressing GFP under the kiss1 gene promoter: Kiss1^hrGFP^ (JAX®, stock 023425) (32), wild type C57B6/J (JAX®; Stock 000664) and the B6.Cg-Lepob/J strain, homozygous mice with an obese spontaneous mutation (*Lep^ob^*, JAX®; Stock 000632). Adult animals used were postnatal (P) age 60-100 days old, unless otherwise specified.

### Ovariectomy and estradiol replacement

To assess the effects of estradiol (E2) on *Slc6a3* gene expression we used ovariectomized (OVX, n=5), OVX + E2 (n=5) and diestrous females (n=4). Females were deeply anesthetized with isoflurane and underwent bilateral OVX. OVX females received steroid replacement via a Silastic capsule containing E2 (1 μg, OVX+E2) or oil (OVX) subcutaneously at the time of surgery. OVX females were perfused 7-14 days following surgery, while OVX+E2 females were perfused two days following E2 replacement. Uterus size was used as control for the treatment. Only OVX mice with uterine weight below 80 mg and OVX+E2 mice with uterine weight above 100mg were used. Both groups were perfused in the morning to avoid time-of-day effects of estradiol feedback.

### Leptin treatment

DAT*^Cre^*;tdTomato adult (P60-70) males and females fasted overnight and prepubertal (P19) males and females fasted for 4h were intraperitoneally (i.p.) injected with leptin (2.5 mg/kg, National Hormone and Peptide Program, Harbor-UCLA Medical Center, CA) or saline (n=5-6 animals/group). Sixty minutes following leptin injection, mice were perfused with PBS and 10% neutral buffered formalin (NBF, Sigma), brains were postfixed for 2h in 20% sucrose in 10% NBF and stored with 20% sucrose in PBS. 30 µm coronal tissue sections 120 µm apart were processed for pSTAT3 immunohistochemistry as described below.

Two cohorts of adult wild type females in diestrus i.p. injected with saline*, Lep^ob^* animals i.p. injected with saline (Lep*^ob^*+ saline) or with leptin (3 mg/kg/day, Lep*^ob^* + leptin group, murine leptin, Preprotech), received the treatment for two days at 9 am and 5 pm and one day at 9 am. One hour after the last saline or leptin injection (at 10 am), females were euthanized by decapitation following anesthesia (isoflurane) and brains were harvested and snap frozen. Coronal frozen sections (16 µm) were collected on a cryostat and stored at -80°C until processing for gene expression.

### In situ hybridization (ISH) with radioisotopes

Adult wild type (WT) male (n=3) and female (n=4) mice were used to study sex differences in *Slc6a3* gene expression. Female mice were also used to determine developmental differences in *Slc6a3* gene expression, *i.e.,* prepubertal (P19, n=7) *vs.* adult (P60-70, n=5) diestrous mice. To assess the effects of nutritional factors in development on *Slc6a3* gene expression we used P20 females from small litters (SL 2-3 pups/litter, n=5 females) or normal litter (NL 7-9 pups/litter, n=4 females). To assess the effect of leptin in *Slc6a3* expression we used diestrous (n=7), Lep*^ob^* + saline (n=5) or Lep*^ob^* + leptin (n=5) injected females. Coronal sections were used for radioactive ISH, using an ^35^S-UTP or ^33^P-UTP labelled *Slc6a3* riboprobe. The following primers were used (exon 10-15 of the *Slc6a3* gene): Forward (5’ ACGTCTTGATCACTGGGCTTGTCGATGAGTT 3’) and reverse (5’ GCATGGATTGGGTGTGAACAGTC 3’) to amplify a 754 base-pair sequence in the *Slc6a3* gene (exons 10-15). A clamp sequence followed by sequences for T7 (CCAAGCCTTCTAATACGACTCACTATAGGGAGA) and T3 (CAGAGATGCAATTAACCCTCACTAAAGGGAGA) promoters were added to the reverse and forward primer sequences, respectively.

Single-labeled ISH was performed on 20 µm fresh frozen or 30 µm fixed brain (120 µm distance) sections mounted onto SuperFrost Excell or Superfrost Gold slides (Fisher Scientific). Fixed sections were subjected to a 10-minute microwave sodium citrate (pH 6) pre-treatment and hybridized overnight at 57 °C with ^35^S-labeled *Slc6a3* riboprobes, as previously described (17, 33). Frozen sections were fixed in ice-cold 10% NBF, treated with 0.25% acetic anhydride and underwent dehydration in ethanol, and hybridized overnight at 57 °C with ^33^P-labeled *Slc6a3* riboprobes. All slides were then incubated in 0.002% RNase A followed by stringency washes in sodium chloride-sodium citrate buffer (SSC). Slides were exposed to film autoradiography (Kodak), for 3-5 days. Slides were dipped in autoradiographic emulsion (Kodak), dried for 3 hours and stored in light-protected boxes at 4 °C for 2-4 weeks. Slides were developed in D-19 developer, dehydrated in ethanol, cleared in xylene, and coverslipped with DPX (Electron Microscopy Sciences). Film images were acquired using a stereoscope (Zeiss). Darkfield 10x images were captured using a digital camera on an AxioImager M2 microscope (Zeiss). ISH signals were quantified using integrated optical density (IOD) in ImageJ software (NIH) using the “freehand” tool to outline the PMv. IOD from the tissue background of the same area was subtracted.

### Fluorescent ISH

We used diestrous WT female mice (n=3) to assess *Slc6a3* and *Lepr* mRNA co-expression by fluorescent ISH. ISH was performed on fresh frozen 16-µm thick cryostat sections at 128-µm resolution (8-series). The ISH was performed following the RNAscope protocol for fresh frozen sections, using Protease III (ACDbio, RNAscope Multiplex Fluorescent Reagent Kit v2). Briefly, slides were dried at 60 °C for 15 min, rinsed in PBS for 5 min, fixed in 10% NBF for 15 min at 4 °C, rinsed in PBS-DEPC 2 times for 3 min, dehydrated through rinses in serial ethanols for 3 min each, and air-dried for 20 min. A hydrophobic barrier was created around each slide using the ImmEdge pen (Vector Laboratories). The slides were then incubated in H_2_O_2_ for 10 min at RT followed by incubation with Protease III for 30 min at 40 °C. ISH was performed using the RNAscope Protease III (ACDBio). Sections were incubated with Mm-Slc6a3-C1 (#315441), and Mm-Lepr-C3 (#402731-C3, labeling all *Lepr* isoforms) RNAscope probes for 2 h at 40°C using the HybEZ Humidifying System (ACDBio). After all incubation steps following the kit’s protocol, slides were incubated in DAPI solution for 30 s at room temperature, and coverslipped using ProLong Gold Antifade Mountant (ThermoFisher Scientific).

Quantification of mRNA coexpression within cells was performed on PMv images acquired with a 40x oil objective on an AxioImager M2 microscope (Zeiss). Based on observed background outside of the area of interest, a threshold for a minimum number of 5 puncta per cell was used to consider a cell positive for expression of that gene. Confocal images were acquired for illustration on a Nikon A1 confocal microscope.

### Electrophysiological recordings

Hypothalamic slices from adult DAT*^Cre^*;tdTomato male (30) and female (31) mice were prepared and the data analyzed as previously described (22). Briefly, mice were decapitated following isoflurane anesthesia, and the entire brain was removed. After removal, the brains were immediately submerged in ice-cold, carbogen-saturated (95% O_2_ and 5% CO_2_) artificial cerebrospinal fluid (ACSF, 126 mM NaCl, 2.8 mM KCl, 26 mM NaHCO_3_, 1.25 mM NaH_2_PO_4_, 1.2 mM MgSO_4_, 5 mM glucose and 2.5 mM CaCl_2_). Coronal sections (250 µM) from hypothalamic blocks were cut on a Leica VT1000S vibratome and incubated in oxygenated ACSF at room temperature for at least 1 hour before the recordings. The slices were transferred to the recording chamber and allowed to equilibrate for 10–20 min. The slices were bathed in oxygenated ACSF (32°C) at a flow rate of ∼2 mL/min. The pipette solution was in some cases modified to include an intracellular dye (Alexa Fluor 488) for whole-cell recording: 120 mM K-gluconate, 10 mM KCl, 10 mM HEPES, 5 mM EGTA, 1 mM CaCl_2_, 1 mM MgCl_2_, 2 mM (Mg)-ATP, and 0.03 mM AlexaFluor 488 hydrazide dye, pH 7.3. Whole-cell patch-clamp recordings were performed on tdTomato-positive neurons anatomically restricted to the PMv. Epifluorescence was briefly used to target the fluorescent cells; at which time the light source was switched to infrared differential interference contrast imaging to obtain the whole-cell recording (Leica DM6000 FS equipped with a fixed stage and a fluorescence digital camera). In current-clamp mode, tdTomato neurons were recorded under zero current injection (I = 0) in whole-cell patch-clamp configuration. The recording electrodes had resistances of 5-7 MΩ when filled with the K-gluconate internal solution. The membrane potential values were compensated to account for the junction potential (-8 mV). In males and females at both ages, the resting membrane potential (RMP) was monitored for at least 10-20 minutes (baseline period) before leptin was administered to the bath. Solutions containing leptin (100 nM) were typically perfused for 15-20 minutes after the baseline period, with a 20-minute washout with ACSF.

### Manipulation of estrous cycles using chemogenetics

Adult virgin DAT-Cre females (n=14) received bilateral injection (50 nL/side) of an adenoassociated virus (AAV) expressing a Cre-dependent hM3Dq-mCherry fusion protein,pAAV8-hSyn-DIO-hM3D(Gq)-mCherry (AAV-hM3Dq, Addgene plasmid # 44361, from Bryan Roth) (34) in the PMv using the following coordinates: Anteroposterior = -5.4 mm (from rostral rhinal vein); mediolateral = -0.52 mm (from sagittal sinus); dorsoventral = -5.4 mm (from dura mater). The stereotaxic protocol is described in detail in a previous study (35). One month after surgery, we started to follow the reproductive cycle of these females collecting daily vaginal smears with saline solution. After a period of adaptation, we continued for 13 days with added DMSO (0.0068%) in water, as a control. Next, we added CNO (5 mg/kg) dissolved in DMSO in the drinking water and followed the cycles for 13 more days. The abundant presence of cornified cells in the smear was considered as estrus/metestrus, a large abundance of leukocytes was considered as diestrus and a large abundance of nucleated cells with some cornified cells present was considered as proestrus. At the end of the experiment, the females received i.p. injection of CNO and 2 h after mice were perfused with PBS and 10% NBF (Sigma), brains were postfixed for 4 h in 20% sucrose in 10% NBF and stored with 20% sucrose in PBS. We collected 30 µm coronal tissue sections 120 µm apart. Sections were cryoprotected and frozen until processed to verify the injection sites for these females.

### Tracing PMv-DAT neuronal projections

DAT-Cre adult females (n=7) received unilateral stereotaxic injections of an AAV expressing a Cre-dependent channelrhodopsin-mCherry fusion protein (AAV8-hSyn-double floxed-hChR2(H134R)-mCherry, UNC Vector Core, from Karl Deisseroth (Addgene 20297), 25-50 nL) in the PMv. One month after the stereotaxic surgery, mice were perfused with 10% NBF and brains were harvested and processed for histology as above. Fixed coronal 30 µm hypothalamic brain sections were processed for immunofluorescence.

### Immunohistochemistry

Fixed frozen tissue sections (30 µm at 120 µm distance) from perfused animals obtained with a freezing microtome (Leica) were rinsed in PBS and blocked with PBS + Triton-X 0.25% (PBT) and 3% normal donkey serum (NDS). Primary antibodies were incubated in PBT + 3% NDS overnight at room temperature. Primary antibodies used were Rabbit dsRed antibody (1:5000, Clontech 632496, RRID:AB_10013483), rat monoclonal anti-mCherry 16D7 (1:5000, Invitrogen M11217, RRID:AB_2536611), rabbit anti-cFOS (1:5000, Millipore ABE457, RRID:AB_2631318), chicken anti-GFP (1:10000, Aves GFP-1010, RRID:AB_2307313), rabbit polyclonal anti-GnRH (1:5,000, Phoenix Pharmaceuticals H-003-57, RRID:AB_572248), and sheep anti-TH (1:5000, Millipore AB1542, RRID:AB_90755). For detection of Fos, endogenous peroxidase was blocked with 0.3% H_2_O_2_ for 30 min before the NDS blocking step. For detection of phosphorylation of signal transducer and activator of transcription 3 (pSTAT3), tissue was pre-treated with 1% H_2_O_2_ and with 1% sodium hydroxide in water and then with 0.3% Glycine before blocking with PBT + NDS 3% as before. Tissue was incubated in primary rabbit anti-pSTAT3 (Tyr705) (D3A7) XP^®^ (1:1,000; Cell Signaling 9145S, RRID:AB_2491009) for 48 h at 4°C. The corresponding secondary fluorescent antibodies were used for detection (1:500, Invitrogen). For the Fos antibody we performed immunoperoxidase detection using a biotinylated-anti rabbit IgG secondary antibody (1:1000, Jackson Immunoresearch), signal amplification with Avidin Biotin Complex (Vectastain^®^ ABC-HRP Kit, 1:500, Vector labs) for 1 h and signal development with diaminobenzidine (DAB, Sigma) 0.05% and 0.01% H_2_O_2_. Floating sections were then mounted on gelatin-coated slides, dried overnight and coverslipped with Fluoromount-G (Invitrogen).

Photomicrographs were acquired using Axio Imager M2 (Carl Zeiss Microscopy). Quantification of pSTAT3, tdTomato and TH positive neurons was performed by an observer unaware of the images’ identity. Dual-labeled tdTomato and pSTAT3 immunoreactive cells were counted in each individual channel and colocalization was considered where pSTAT3 immunoreactivity (-ir) was clearly nuclear in tdTomato-positive cells. Two sections at the mid-PMv level were counted (∼Bregma: -2.46 mm). No correction for double counting was performed because sections were 120μm apart. tdTomato fiber density in the AVPV/PeN was quantified using IOD in ImageJ software (NIH) on both sides of the ventricle in a representative section for each region. An elongated rectangle of the same size for all animals was used as region of interest, placed in contact with the ventricle wall to cover a representative area over the TH-expressing cells.

Confocal microscopy images were acquired and analyzed using a Nikon A1 microscope and a Nikon N-SIM + A1R microscope with a resonance scanner.

### Data analysis

Data are expressed and represented as mean ± SEM. When data did not fit a normal distribution or did not have equal variances, they were transformed to fit a normal distribution and re-analyzed. Unpaired two-tailed Student’s t test was used for comparison between two groups. For comparison between three groups, one-way ANOVA was used followed by Tukey’s *post-hoc* multiple comparison test. For pSTAT3 and %pSTAT3/tdTomato cells, a two-way ANOVA was used with age and sex as factors. Correlation was assessed between body weight and *Slc6a3* gene expression using Pearson R correlation coefficient. A P value less than 0.05 was considered significant. Data were organized and calculated in Excel software (Microsoft, inc.). Statistical analyses and graphs were performed using GraphPad Prism v.9.5 (GraphPad software, inc.). Zen Blue 3.7 software (Carl Zeiss Microscopy GmBH) was used to acquire and process epifluorescence images. NIS-elements software (Nikon) was used to acquire and process confocal microscope images. Photoshop 2024 (Adobe, inc.) was used to integrate graphs and digital images into figures. The graphical abstract was prepared using BioRender. Only brightness, contrast, and levels were modified to improve data visualization in the figures.

## Results

### *Slc6a3* mRNA expression in the PMv is sexually dimorphic and higher in prepubertal females

The *Slc6a3* (DAT) gene is expressed in the PMv of male and female mice (24, 25). To evaluate potential sexual dimorphism or postnatal developmental changes, we assessed *Slc6a3* gene expression in the PMv in adult males and females, and in prepubertal and adult females. PMv *Slc6a3* mRNA levels were higher in diestrous females compared to male mice (n=3 female, n=4 male, unpaired t-test p=0.032 Figure 1A, B, D), and higher in prepubertal females compared to diestrous mice (n=7 prepubertal, n=5 diestrous, unpaired t-test p=0.013 Figure 1B, C, E).

**Figure 1.**
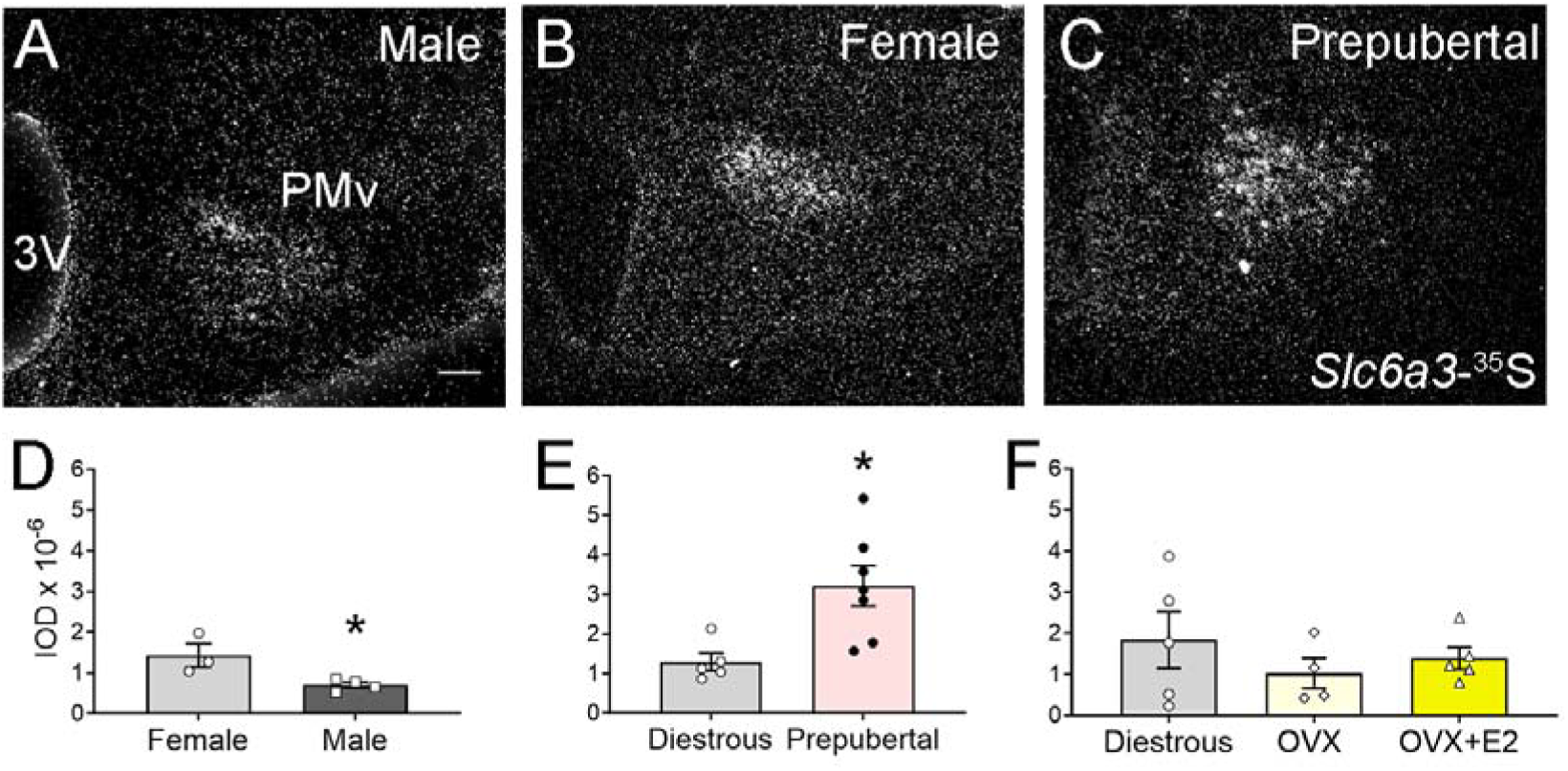
Ventral premammillary nucleus (PMv) *Slc6a3* gene expression varies with sex and development. **A-C.** Darkfield images showing the *Slc6a3*-^35^S hybridization signal (silver grains) in the PMv of adult male, a diestrous female and a prepubertal female, respectively. **D-F.** Graphs showing the quantification of the *Slc6a3* hybridization signal in adult male *vs*. diestrous females, in prepubertal *vs*. diestrous females and in diestrous *vs*. ovariectomized (OVX) females and OVX females supplemented with estradiol (E2). All data shown are average ± SEM. * p<0.05. Scale bar = 100 µm.

To assess if the reduction of PMv *Slc6a3* mRNA in adult female mice is a result of increasing circulating estradiol (E2) during the pubertal transition, hypothalamic sections from diestrous, ovariectomized (OVX), and OVX + E2 mice were analyzed. We found no differences between these groups (n=5 diestrous and OVX+E2, n=4 OVX, one-way ANOVA, *p* = 0.54, Figure 1F).

### PMV^DAT^ neurons show heterogenous responses to leptin

We used fluorescent *in situ* hybridization (ISH) to assess transcript coexpression in adult females in diestrus (n=3). Virtually all *Slc6a3* neurons in the PMv coexpressed *Lepr* mRNA (93.6 % ± 2.1), whereas about half of PMv *Lepr* neurons coexpressed *Slc6a3* mRNA (58.6 % ± 4.1, Figure 2A-C).

**Figure 2.**
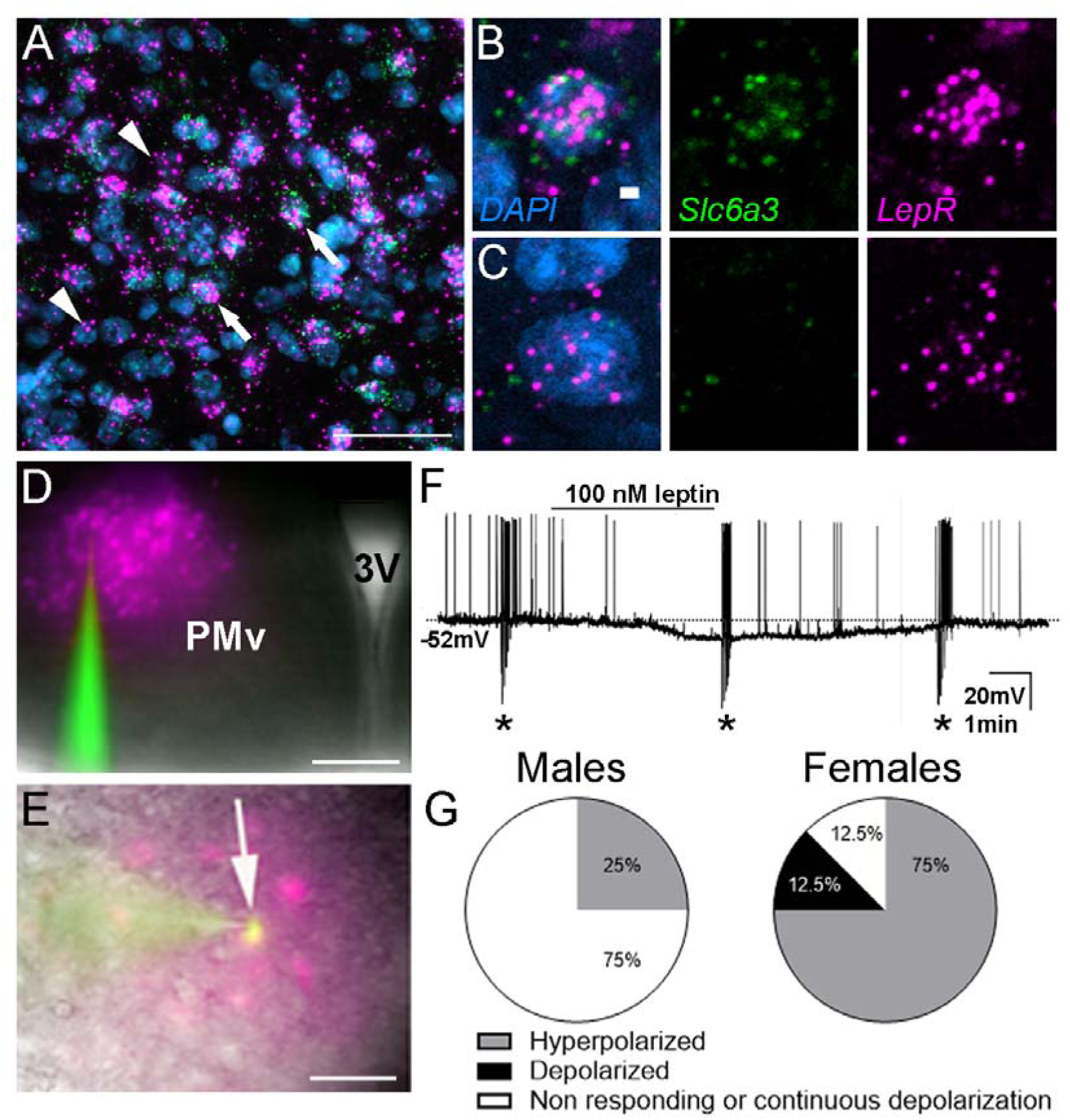
A subpopulation of leptin receptor (*Lepr*) expressing neurons in the ventral premammillary nucleus (PMv) co-expresses dopamine transporter (*Slc6a3)*. **A.** Fluorescent image showing representative fluorescent *in situ* hybridization depicting the colocalization of *Lepr* (magenta) and *Slc6a3* (green) in the PMv of a diestrous female. Arrows point to cells co-expressing *Lepr* and *Slc6a3* mRNA. Arrowheads point to cells expressing only *Lepr*, but not *Slc6a3*, mRNA. Blue = DAPI. **B-C.** Higher magnification of individual cells depicted in A that co-express *Slc6a3* and *Lepr* mRNA (B), and of a cell that expresses only *Lepr* mRNA (C). **D.** Fluorescent image showing the PMv in the brain slices, recognized by tdTomato expression in DAT*^Cre^* neurons. **E.** Merged image showing the colocalization between a recorded DAT*^Cre^*;tdTomato neuron (magenta) and the AF488 dye (green), dialyzed during the recording. **F.** Representative current-clamp recording demonstrating leptin (100 nM) induced hyperpolarization in a subset of DAT-Cre tdTomato neurons of a male mouse. The dashed line indicates resting membrane potential (-52 mV). Asterisks indicate square pulse current injections to assess input and access resistance. **G.** Pie charts representing the percentage of neurons that hyperpolarized, depolarized or did not respond to 100nM leptin in adult males (N=16) and in adult diestrous females (N=8). Scale bars: A and E = 50 µm, B-C: 2 µm, D= 400 µm.

To investigate the effect of leptin on the membrane excitability of PMv^DAT^ neurons, we performed current clamp recordings of DAT*^Cre^*;tdTomato neurons. In males, the average RMP of recorded neurons was -52.8 ± 2.0 mV (range from -62 to -38 mV, 16 cells from 8 mice). In a separate cohort of females, the RMP was -57.2 ± 5.7 mV (range from -62 to -51 mV, 8 cells from 6 mice). We found that bath application of 100 nM leptin hyperpolarized 25% of the recorded neurons of male mice (4/16 cells Figure 2D-G). The RMP of the remaining 75% of the recorded cells was unchanged or showed a continuous depolarizing trend and were removed from the analysis. In females in diestrus, ∼75% (6/8) of recorded cells hyperpolarized in response to bath application of 100nM leptin (Figure 2G). Two PMv^DAT^ neurons showed a depolarizing response, and one showed continuous depolarization and was removed from the analysis (Figure 2E). The leptin-associated hyperpolarization of PMv^DAT^ cells was of similar amplitude in both sexes: -7.8 ± 0.8 mV in males, and -6.3 ± 2.0 mV in females.

Although most (> 90%) of *Slc6a3* expressing neurons express *Lepr*, only a subpopulation exhibits a response, including both de- and hyperpolarization, in electrical properties to the hormone.

### Long-term activation of PMv^DAT^ neurons does not alter estrous cycles in adult virgin females

Given the higher percentage of female’s DAT*^Cre^*;tdTomato neurons that are hyperpolarized by leptin, we decided to investigate if long-term activation of these neurons might impair the reproductive cycle of the adult females. We stereotaxically injected the AAV-hM3Dq virus bilaterally in the PMv of 14 DAT*^Cre^*;tdTomato virgin females. Twelve females had bilateral injections and one had a unilateral injection centered in the PMv, defined by the expression of mCherry and Fos immunoreactivity that indicates they had been activated by CNO (Figure 3A-B). Two animals had missed injections with not Cherry expression observed in either PMv, and lack of Fos confirmed in the PMv of these animals (Figure 3C). Of the twelve animals with bilateral injection, eleven showed regular cycling (at least two complete cycles) during the control (DMSO) period and were used for the analyses (Figure 3D). The cycles of the bilaterally injected females were not altered by the CNO when compared to the DMSO exposure (paired t-test DMSO *vs*. CNO, days in estrus/metestrus, p=0.59; days in diestrus, p=0.68; cycle length, p=0.69, n=11, Figure 3E-G). We paid special attention to the potential virus spread to a nearby population of *Slc6a3* expressing cells, the tubero-infundibular dopamine (TIDA) neurons in the arcuate nucleus (Arc), (24). Six mice showed some viral contamination of TIDA neurons, but no differences in cyclicity or cycle length were noticed when these animals were removed from the analysis. These results suggest that these cells have no effect on female cyclicity.

**Figure 3.**
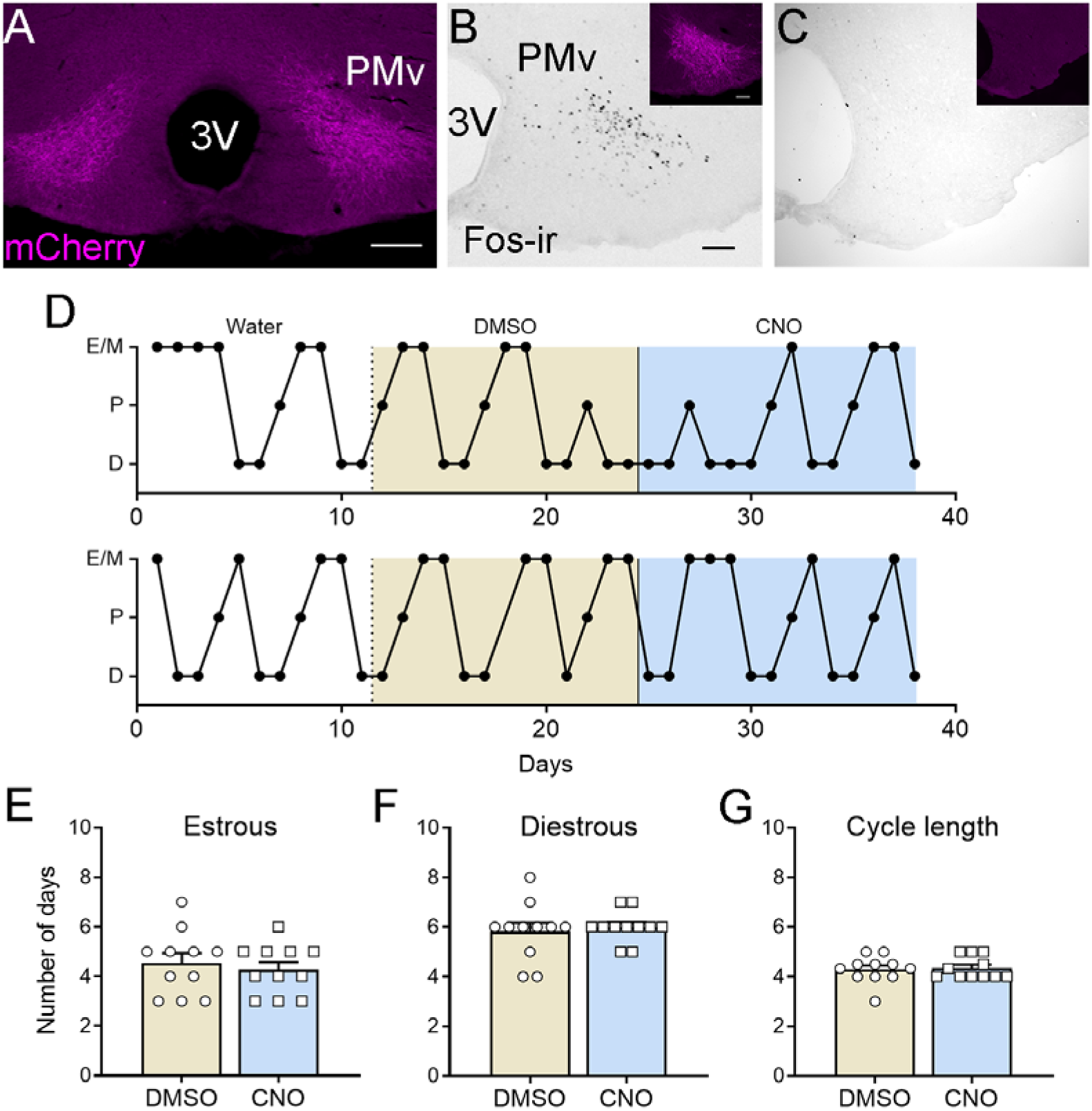
Continuous activation of dopamine-transporter neurons in the ventral premammillary nucleus (PMv^DAT^) does not alter estrous cycles in adult DAT^Cre^ female mice. **A.** Representative low-magnification fluorescent image of bilateral injections of adenoassociated virus (AAV) expressing Cre dependent hM3Dq-mCherry targeted to the PMv of a female DAT^Cre^ mouse. **B.** High magnification image showing Fos immunoreactivity (Fos-ir) in one PMv side following an intraperitoneal injection of clozapine-N-oxide (CNO) and corresponding fluorescent image of the PMv showing mCherry immunofluorescence. **C.** High magnification image showing the lack of Fos-ir in one PMv side of a missed AAV injection, following an intraperitoneal injection of clozapine-N-oxide (CNO) and corresponding fluorescent image of the PMv showing the lack of mCherry immunofluorescence. **D.** Representative estrous cycles of two females with AAV injections centered in the PMv before treatment (drinking water), during the treatment with DMSO (vehicle) and CNO in drinking water. E/M: estrus/metestrus; P: Proestrus; D: Diestrus. **E-G.** Graphs showing the number of days spent in estrus/metestrus, diestrus and the cycle length (number of days) in the DAT^Cre^ females with bilateral PMv AAV-hM3Dq injections during the DMSO and the CNO treatment. Data are average ± SEM. Scale bars: A = 200 µm, B, C = 100 µm

### Prepubertal PMv^DAT^ neurons respond to leptin and show distinct membrane properties compared to adult females

Due to increased expression of *Slc6a3* in prepubertal females, we explored the functional response of PMv^DAT^ neurons to exogenous leptin. Adult and prepubertal DAT*^Cre^*;tdTomato mice received an i.p. injection of leptin, and one hour after, colocalization of pSTAT3-ir in tdTomato neurons was quantified. No differences were observed in the number of pSTAT3-ir cells with age or sex in leptin treated mice (n = 3-5; Two-way ANOVA, p=0.28 for Sex; p=0.79 for Age; Figure 4A-G). Virtually no pSTAT3-ir was observed in the PMv of saline treated mice (n=5 per group). As expected from the *Slc6a3* and *Lepr* coexpression data, 95.3 – 99.3 % of tdTomato neurons in the PMv colocalized with pSTAT3-ir in adults of both sexes (Figure 4H). Similar colocalization was observed in prepubertal mice of both sexes (96.4 – 98.9 %, Figure 4H). About 30% of PMv pSTAT3-ir neurons colocalized with tdTomato in females (25.9 ± 2.8% prepubertal, and 33 ± 2.4% in diestrous, p=0.14) and males (27.8 ± 1.8% prepubertal, and 35.6 ± 4.8% in adults, p=0.18).

**Figure 4.**
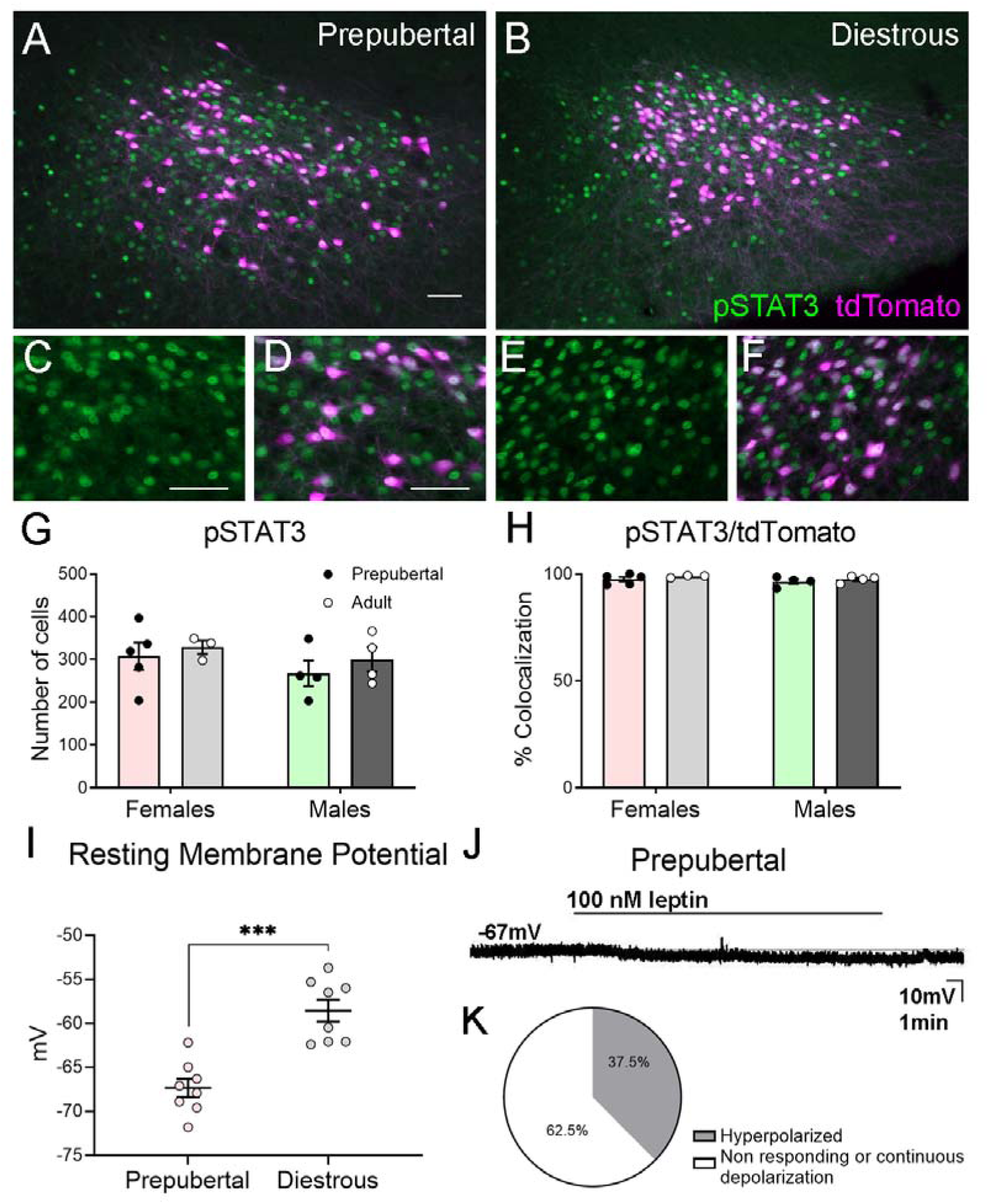
Prepubertal dopamine-transporter neurons in the ventral premammillary nucleus (PMv^DAT^) are responsive to leptin but are more hyperpolarized than in adult females. **A, B.** Fluorescent images showing the colocalization of pSTAT3 (green) and tdTomato (magenta) immunoreactivities in the PMv of prepubertal and adult diestrous DAT*^Cre^*;tdTomato females 1h after a 2.5 mg/kg intraperitoneal leptin injection. **C-F**: High magnification images of the images depicted in A (C, D) and B (E, F). **G, H.** Graphs showing the number of pSTAT3 neurons per section (G) and the percentage of tdTomato cells expressing pSTAT3 (H) in the PMv of prepubertal and adult DAT*^Cre^*;tdTomato female mice. Prepubertal (n=5) and diestrous (n= 3) females, prepubertal and adult males (n=4 each). **I.** Graph showing the resting membrane potential of individual cells from prepubertal and adult females (n=8 each). *** p<0.001. **J.** Representative current-clamp recording demonstrating leptin (100 nM) - induced hyperpolarization in a DAT-Cre tdTomato neuron of a female prepubertal mouse. The dashed line indicates resting membrane potential (-67 mV). **K.** Pie chart representing the percentage of neurons that hyperpolarized or did not show a change in membrane potential to 100nM of leptin in prepubertal females (N=8). All data shown are average ± SEM. Scales in A-B = 50 µm. C-F = 20 µm.

When examined by electrophysiology, the PMv^DAT^ neurons of prepubertal (unweaned) female mice revealed heterogeneous properties. Interestingly, the RMP of prepubertal PMv^DAT^ neurons was more hyperpolarized compared to adult diestrous females (n=8 prepubertal and n=8 diestrous, unpaired t-test, p <0.0001, Figure 4I). In response to leptin treatment, three out of eight cells (37.5%) from three mice showed no RMP change and another three out of eight cells (37.5%) responded by hyperpolarization (Figure 4J, K). Two out of eight cells (25%) depolarized after acute leptin, but none of them recovered after washout. Two recorded cells showed continuous depolarization and were removed from the analysis. The hyperpolarized cells showed a -6.3 ± 2.1 mV change in the RMP after treatment.

Our findings indicate that the PMv^DAT^ neuron population from prepubertal (unweaned) female mice is in a less excitable state compared to adult mice.

### Postnatal overnutrition increases *Slc6a3* mRNA expression in the PMv of prepubertal females

We next assessed if the expression of *Slc6a3* mRNA is altered in the PMv of leptin deficient *Lep^ob^* infertile female mice, which remain in a prepubertal state. We employed a paradigm of leptin treatment and pubertal progression in which *Lep^ob^* females were injected with saline or leptin twice a day for 2 ½ days are compared to age-matched wild type diestrous females (16, 36). As expected, the leptin-treated *Lep^ob^* mice showed a significant decrease in body weight and displayed signs of pubertal progression (vaginal opening) following the leptin treatment. We found that *Slc6a3* mRNA expression is ∼40% lower in non-treated *Lep^ob^* females, compared to wild type females in diestrus (n=5-7; p=0.0025 one-way ANOVA, Tukey’s *post-hoc*, p=0.005 Figure 5A). The short-duration leptin treatment regimen was sufficient to induce a ∼ 40% increase in *Slc6a3* mRNA expression in the PMv of *Lep^ob^* mice (Tukey’s *post-hoc*, p=0.005, Figure 5A), concomitant with the first signs of puberty onset.

**Figure 5.**
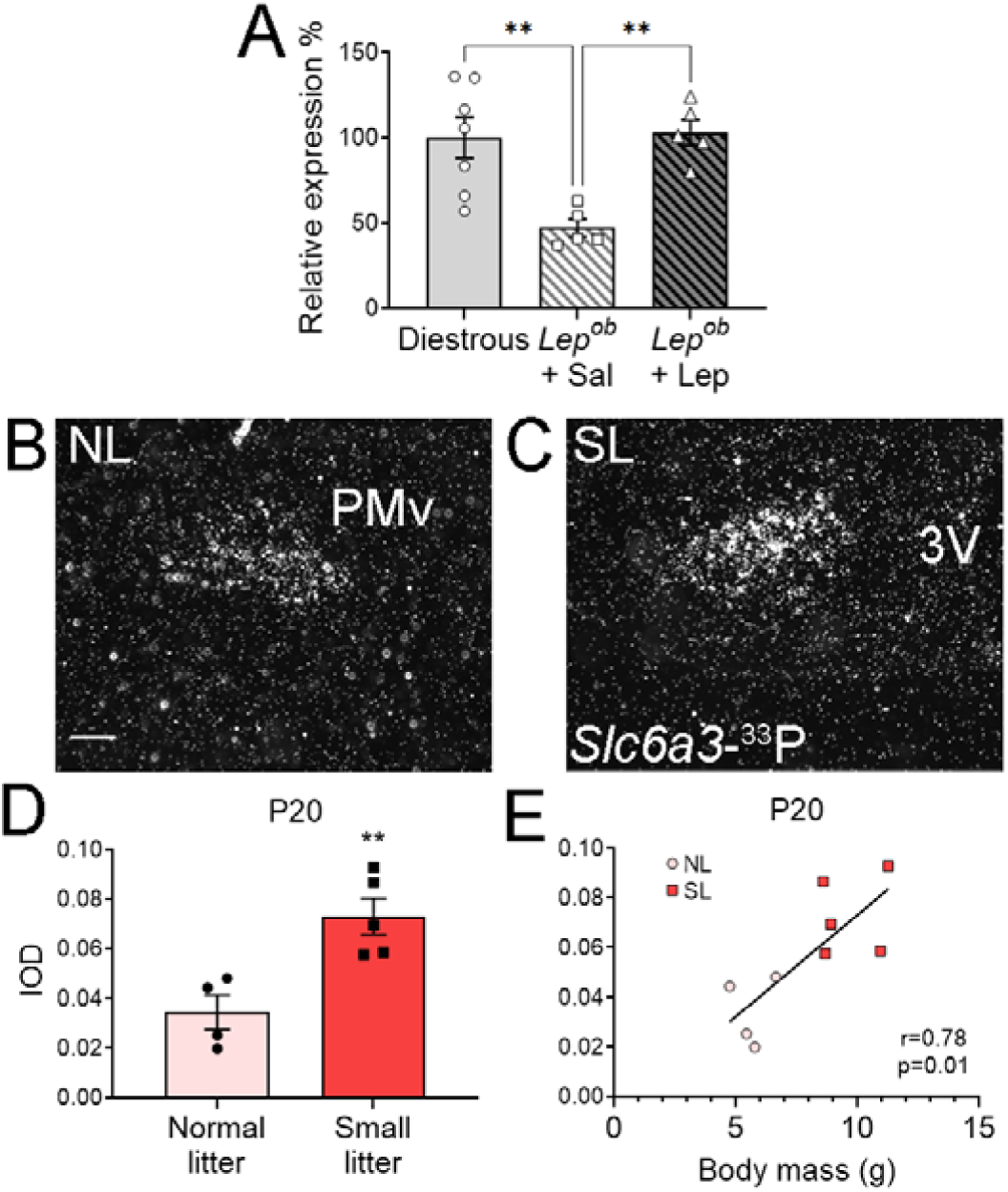
*Slc6a3* (DAT) gene expression in the ventral premammillary nucleus (PMv) of prepubertal mice is affected by nutrition and leptin. **A.** Graph showing the quantification of *Slc6a3*-^33^P hybridization signal as relative expression (%) in diestrous wild type *vs*. prepubertal *Lep^ob^* females injected with saline or with leptin. n=7 for diestrous and for *Lep^ob^* + saline, n = 5 for *Lep^ob^* + leptin. **B-C.** Dark-field images showing the *Slc6a3*-^33^P hybridization signal (silver grains) in the PMv of prepubertal females (P20) from normal litter (NL) size (B) and from small litter (SL) size (C). **D.** Graph showing the quantification of the *Slc6a3* hybridization signal (integrated optic density) in P20 females from normal and small size litters. n=4 animals for normal litter and n = 5 animals for small litter. **E.** Graph showing the correlation between female body mass and the *Slc6a3* hybridization signal (IOD) in P20 females from normal and small size litters. Data shown are average ± SEM. ** p<0.01. Scale bar in B = 100 µm.

Given the complex phenotype of the *Lep^ob^* mouse (37), we employed a paradigm of postnatal overnutrition, which leads to high leptin levels and early puberty (38, 39). We compared *Slc6a3* mRNA expression in the PMv of females raised in normal (NL, n=4 females) versus small (SL, n=5 females) litter sizes. Body weight was higher in SL offspring as compared to NL (9.67 ± 0.59 g in SL *vs*. 5.65 ± 0.39 g in NL, unpaired t-test p=0.001). *Slc6a3* mRNA levels were higher in the SL than in those in NL (p=0.007, Figure 5B-D) and were strongly correlated to body weight (Pearson r=0.78; p=0.01, Figure 5E).

### PMv^DAT^ neurons project to kisspeptin AVPV/PeN neurons

To assess if PMv^DAT^ neurons are part of the circuitry regulating pubertal development, DAT*^Cre^*;*Kiss1^hrGFP^*females were unilaterally injected with a Cre-dependent AAV expressing a channelrhodopsin-mCherry fusion protein (AAV-ChR2-mCherry) into the PMv (n=7). Abundant mCherry-ir neurons were observed within the PMv in correctly targeted animals (n=6). Mice showing virus spread to nearby DAT expressing populations were removed from the analysis (n=2 were analyzed, Figure 6A). In accordance with previous studies focused on PMv projections (41, 42), dense mCherry-ir fibers were found in several hypothalamic regions including the AVPV (Figure 6B), the periventricular nucleus (PeN) the medial preoptic area (MPA, Figure 6C), and the ventrolateral subdivision of the ventromedial hypothalamus (VMHvl, not shown). Most notably, very sparse innervation of the Arc was observed (Figure 6D). No mCherry-ir projections were observed nearby or in contact with GnRH cell bodies in the medial septum (MS) or MPA (not shown).

**Figure 6.**
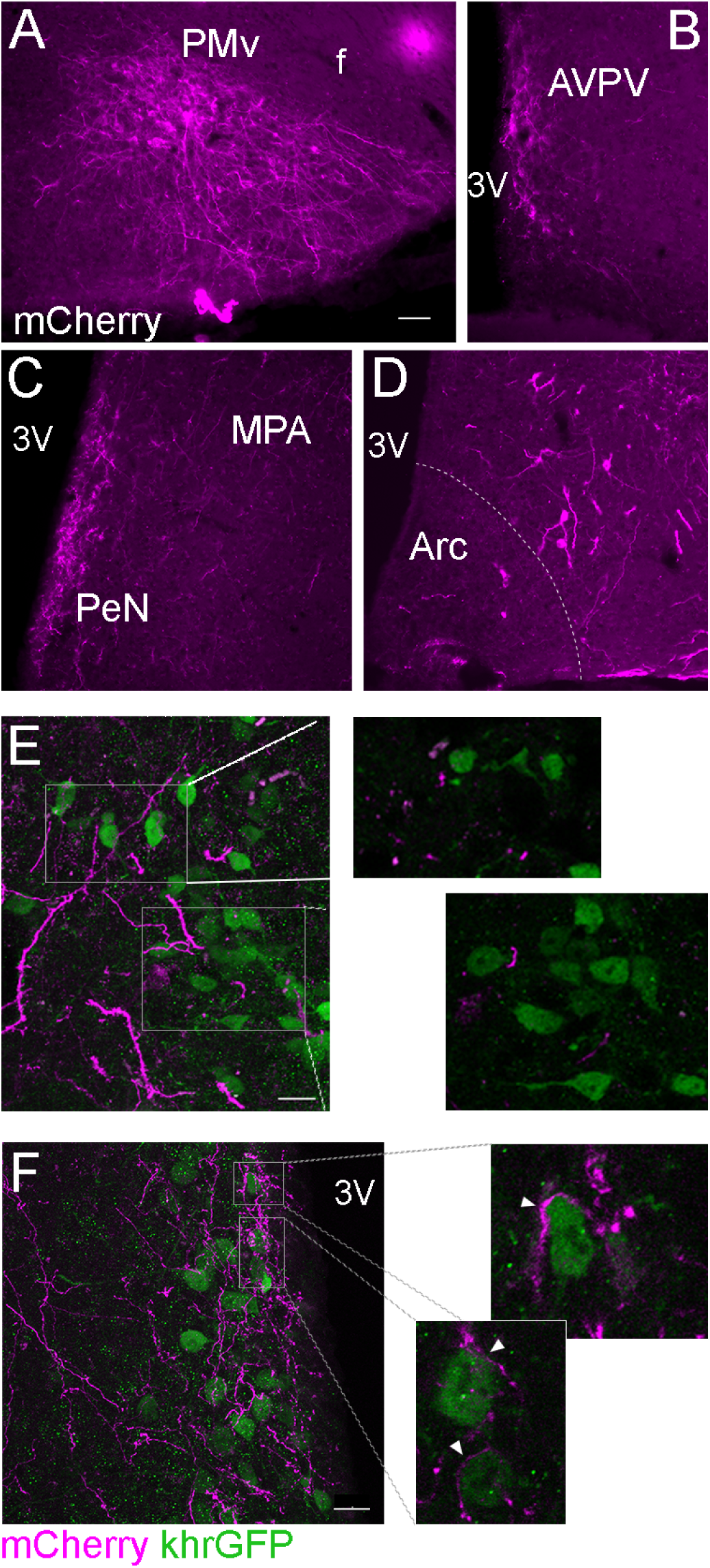
Dopamine-transporter neurons in the ventral premammillary nucleus (PMv^DAT^) project to the AVPV and PeN and contact kisspeptin neurons. **A-D**. Fluorescent image showing ChR2-mCherry signal at the site of the injection in the PMv A, and the projections to the Anteroventral periventricular nucleus (AVPV, B), the Periventricular nucleus (PeN) and the Medial preoptic area (MPA, C), and presence of few projections in the arcuate nucleus (Arc, D). **E.** Confocal fluorescent maximum intensity projection image showing the lack of mCherry (magenta) projections close to kisspeptin (*kiss1*hrGFP, khrGFP) neurons (green) in the Arc. Insets are higher magnification single z plane images showing detail of mCherry signal in the external part of the dorsal Arc and the lack of contacts to khrGFP neurons in this area. **F.** Confocal fluorescent maximum intensity projection image showing mCherry signal (magenta) in the AVPV/PeN and the intense interaction of these projections to khrGFP neurons (green) in this area. are higher magnification single z plane images showing detail of the close contacts between these two populations (arrowheads). f: fornix; 3V: Third ventricle. Scale bar A-D = 50 µm. E-F = 20 µm.

To explore a possible interaction of PMv^DAT^ neurons with kisspeptin, we analyzed mCherry-ir fibers in proximity to *Kiss1^hrGFP^*cells using confocal microscopy. As expected, due to the low innervation of the Arc, kisspeptin/neurokinin 3/dynorphin (KNDy) neurons did not receive close appositions from PMv^DAT^ neurons (Figure 6E). In contrast, dense mCherry innervation of kisspeptin cells was observed in the AVPV and PeN region (AVPV/PeN, a.k.a. rostral periventricular area of the third ventricle, Figure 6F) of the adult female mouse.

### PMv^DAT^ innervation of AVPV/PeN is established during the pubertal transition

The AVPV/PeN area contains a sexually dimorphic population of dopaminergic TH and kisspeptin cells, both denser in females. About 50-90% of the *Kiss1* cells express TH in mice (40–43), and kisspeptin expression in this region increases during the pubertal transition (40, 44). In DAT*^Cre^*;tdTomato mice, we found that AVPV/PeN TH neurons do not express tdTomato. DAT*^Cre^*;tdTomato fiber density was about three times higher in the AVPV (p=0.006, Figure 7A, B and E), and about ten times higher in the PeN of adult *vs*. prepubertal females (p=0.003, Figure 7C, D and E). As observed for *Kiss1*, the number of neurons expressing TH was higher in both the AVPV and the PeN of adult diestrous female mice compared to prepubertal females (p<0.0001 in the AVPV; p<0.0001 in the PeN, Figure 7A-D, F). DAT*^Cre^*;tdTomato terminals were in close apposition to TH neurons and fibers in the AVPV/PeN of adult female mice (Figure 7G-H).

**Figure 7.**
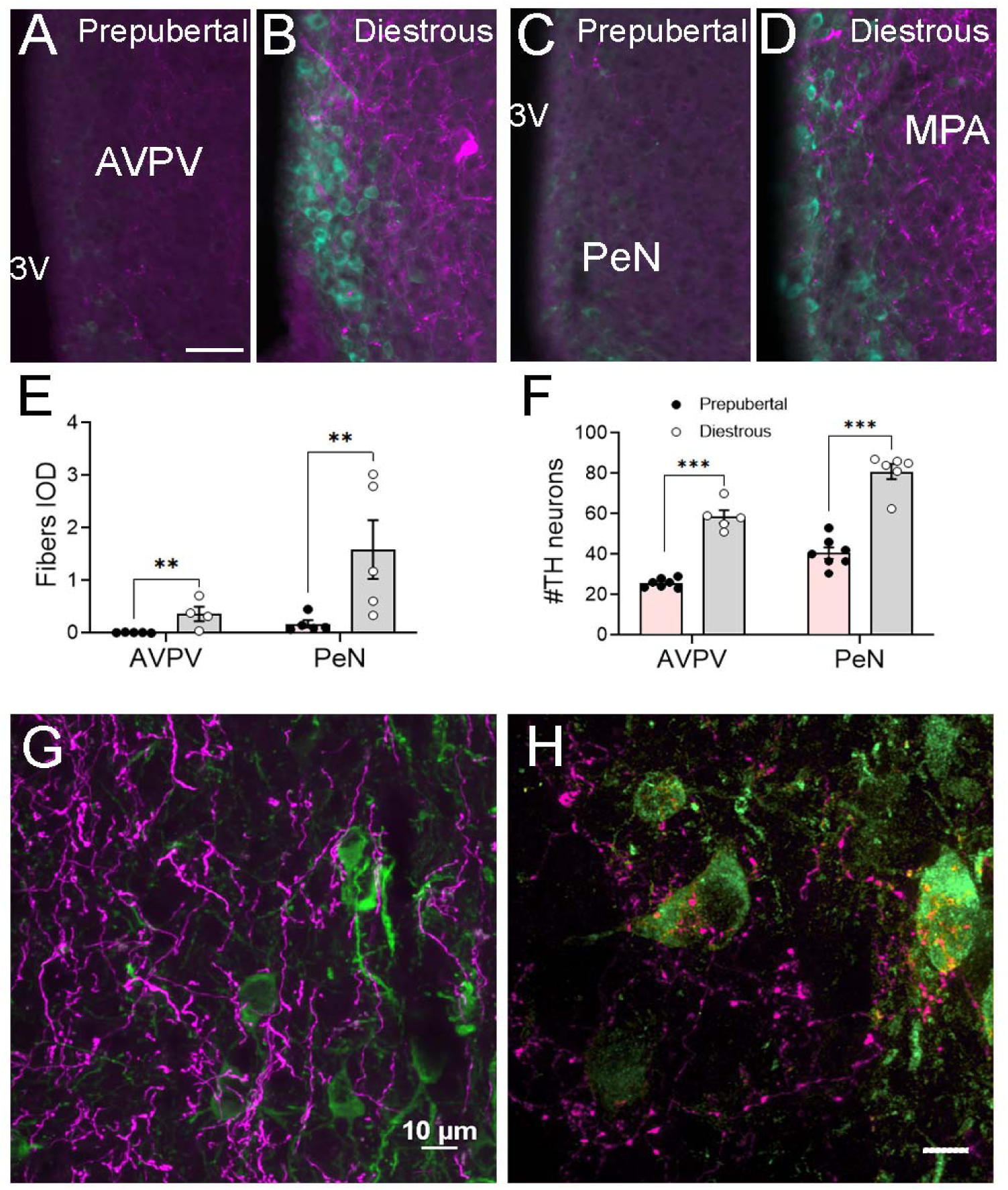
DAT^Cre^;tdTomato projections to dopaminergic cells in the anteroventral periventricular and periventricular nuclei (AVPV/PeN) arise after puberty. **A-D**. Fluorescent images showing tdTomato (magenta) and tyrosine hydroxylase (TH) immunoreactivity (green) in prepubertal (A, C) and in diestrous (B, D) DAT^Cre^;tdTomato females. **E.** Graph showing the quantification of the tdTomato fiber density (Integrated optical density, IOD) in prepubertal *vs*. adult diestrous females in the AVPV and the PeN areas. **F.** Graph showing the number of TH neurons per section in the AVPV and in the PeN in prepubertal *vs*. diestrous female mice. **G.** Confocal fluorescent maximum intensity projection image showing tdTomato (magenta) and TH (green) cells and fibers in the lateral region of the AVPV/PeN in an adult diestrous female. **H.** Higher magnification confocal fluorescent maximum intensity projection image of a different field from G., showing tdTomato fibers in the PeN (magenta) and the intense interaction of these projections to TH neurons (green) in this area. All data shown are average ± SEM. ** p<0.01, *** p<0.001. Scale bars in A-D = 50 µm. G = 10 µm. H = 5 µm.

## Discussion

The present study revealed a novel subset of leptin responsive cells within the PMv that show dynamic regulation of the *Slc6a3* (DAT) mRNA during puberty and specific projections to hypothalamic sites and neurons involved in puberty and reproductive control. The overall PMv^Leprb^ population is key in the metabolic regulation of puberty and fertility, whereas the PMv^DAT^ neurons were previously shown to be involved in aggression, social and maternal behaviour (25, 26, 28, 29). Our findings indicate that the PMv^DAT^ population represents a discrete subset of PMv^LepRb^ neurons and a novel candidate for mediating nutritional modulation of reproduction and pubertal development.

Within the PMv, leptin-induced pSTAT3 was observed in all DAT-expressing cells. However, acute leptin exposure had heterogenous sex- and age-dependent effects on the excitability of DAT^Cre^ neurons. The lack of electrophysiological response of a subpopulation suggests that these neurons are responsive to leptin in ways not tested in this study, *e.g.* by means of transcriptional regulation. Acute leptin action in the PMv*^Leprb^* population is also heterogenous, inducing the depolarization of 75% of these cells, via a putative transient receptor potential channel (TRPC) channel, and hyperpolarizing 25% cells, via activation of a putative Katp channel, while no cells were found to be unresponsive to the treatment (22). Importantly, the RMP of prepubertal females was more hyperpolarized than that of adult females suggesting that these cells may increase in excitability with maturation and only assume a more active role within the circuit after puberty. PMv^DAT^ cells are responsive to prolactin and oxytocin (29, 45) and the influence of these hormones may contribute to switching these cells from a quiescent to an excitable state in their role in maternal aggression (29). Further studies are needed to determine the acute and chronic effects of leptin on the electrical properties of PMv^DAT^ neurons in distinct developmental and physiological states, as well as the mechanisms associated with changes in intrinsic physiological properties of PMv^DAT^ neurons from prepubertal to adults.

PMv^DAT^ cells have been described as a non-dopaminergic population (25, 27). These cells are an active glutamatergic population and send excitatory inputs to projection sites, in particular the VMHvl (25, 26). Several studies, including ours (not shown) have observed a lack of TH expression in this population (27, 46). Only one study has shown an enrichment in TH using Ribotag mice, but to a much lower extent than classical midbrain dopaminergic cells (25). Still, PMv^DAT^ neurons express other elements of the dopamine/monoamine regulation pathway, such as *Gucy2c*, *Aadc* and *Vmat2* (25, 47). In shrews, dopamine and serotonin have been detected in the PMv after L-DOPA and 5-HTP treatment, respectively (47). However, in mice studies using fast-scan cyclic voltammetry have suggested that PMv^DAT^ neurons do not produce dopamine, even when supplemented with L-DOPA (25). Thus, a significant role for dopamine release from these neurons is viewed as unlikely. Still, here we have discerned differences in the regulation of the dopamine transporter that merit further attention.

*Slc6a3* mRNA expression in the PMv was higher in females than in males, and showed a developmental decrease after puberty, suggesting a regulation during sexual maturation. Removal of the ovaries or changes in estrogen levels did not affect *Slc6a3* mRNA expression, making estrogen an unlikely regulator during the pubertal shift. Leptin is critical for puberty and fertility. Here, overnourished prepubertal animals from SL showed increased level of *Slc6a3* mRNA expression in PMv, correlated with individual’s body mass, although other factors, such as altered sex distribution in the mostly-females SL could also affect gene expression. Furthermore, the rescue of *Slc6a3* mRNA levels observed in leptin-treated *Lep^ob^* animals suggests that leptin has an active role in increasing *Slc6a3* gene expression during pubertal transition. Whether other hormones (*e.g.* prolactin, oxytocin), different physiological states or social behaviors affect PMv *Slc6a3* expression is unknown.

Leptin regulation of dopaminergic neurotransmission in the midbrain (substantia nigra and ventral tegmental area) is involved in motivation for food rewards and locomotion (48–50). Leptin specifically regulates dopamine-related genes in these populations (51) and their action in the nucleus accumbens (52). It is important to note though that most studies investigating the role of PMv neurons have used DAT-Cre mice as a strategy to assess the PMv’s neuronal function and circuitry, not DAT expression and function. More studies are needed to reveal the role of dopamine-related genes in the PMv, but our findings suggest that *Slc6a3* and its regulation by leptin have a role in pubertal maturation. DAT is mostly found in presynaptic terminals (53); so, this role might be relevant in PMv projection sites, perhaps regulating the neurotransmission of PMv neurons.

The PMv^DAT^ neurons project to a subset of brain sites innervated by the PMv^Leprb^ population. The PMv^Leprb^ population innervates VMHvl, and key neuronal populations in the control of reproduction, namely, GnRH neurons in the OVLT, MPA and medial septum areas, KNDy neurons in the Arc, and kisspeptin cells in the AVPV/PeN (16, 18, 54). As reported before, PMv^DAT^ neurons project to the VMHvl and the supramammillary nucleus (25, 26). In the MPA, we observed little density of axons. However, we observed a very dense collection of terminals in the AVPV/PeN. Differences with previous studies might arise from the use of a different protein marker (ChR2 *vs*. synaptophysin). Of the reproductive populations targeted by PMv^Leprb^ neurons, the PMv^DAT^ neurons seem to specifically target the kisspeptin population in the AVPV/PeN. These results show that the PMv^DAT^ subpopulation target a group of neurons essential for pubertal development in females, supporting a function in puberty.

In adults, AVPV/PeN kisspeptin neurons mediate the positive feedback action of estradiol on LH surge that precedes ovulation on the afternoon of proestrus (55). We recently showed that acute activation of the PMv^Leprb^ population in females leads to LH release (20). However, chronic chemogenetic activation of PMv*^DAT^* cells had no effect on estrous cycles (present study). Alterations of the LH surge in proestrus have been observed in the absence of effects on estrous cycle progression (56). Therefore, we cannot discard any effects on the LH surge, which can be disrupted after PMv lesions in rats (4, 57). The lack of direct PMv*^DAT^* projections to Arc KNDy or GnRH neurons, the dynamic changes in *Slc6a3* expression and neuronal properties, the dense projections to the AVPV/PeN, and the AVPV/PeN’s intense chemical remodelling during puberty maturation, suggest that the PMv^DAT^ population could undergo a functional switch during pubertal maturation. We hypothesize that the PMv^DAT^ population could play an important role in the regulation of sexual maturation and later reproductive function, integrating the nutritional state from leptin signal into the AVPV/PeN.

Similar to gene expression, the innervation of the AVPV/PeN from DAT^Cre^ neurons was dynamically regulated during female puberty. In adult females DAT projections densely englobed TH neurons, a trait absent in prepubertal females. This timing probably reflects an increase in *Slc6a3*-driven *Cre* expression and a delay to observe tdTomato - ir. The AVPV is a sexually dimorphic population, with higher cell abundance in the female (58). Similar to kisspeptin neurons, the TH population in the AVPV/PeN is sexually dimorphic, with more cells present in females (59, 60).

Notably, TH expression in the AVPV/PeN was much lower in prepubertal females than in the adults, a similar developmental time as kisspeptin appearance (61) and the increase of DAT terminals in the AVPV/PeN. This is significant because the AVPV/PeN region is one of the classical dopaminergic regions of the hypothalamus, also known as A15 area (62). The A15 TH neurons do not express DAT (46, 63, 64). Prototypical dopaminergic neurons (*i.e.,* in the midbrain), co-express TH and DAT, so that dopamine is recycled by DAT at the presynaptic terminal. The significance of our current finding is puzzling. However, a recent report provided evidence that DAT plays an integral role in managing the dopaminergic micro-circuitry within the ARH TIDA neurons (65), suggesting that PMv^DAT^ could participate in the AVPV/PeN dopamine microcircuitry, potentially via expression of Aromatic l-aminoacid decarboxylase (AADC) (25, 47). We therefore speculate that the presence of DAT at the PMv presynaptic terminal plays a role in regulating dopaminergic tone in the AVPV/PeN microcircuits and female reproductive function. In addition, PMv^DAT^ neurons likely act on AVPV cells via glutamatergic signaling. AVPV TH neurons play a role in maternal behavior and intermale aggression (66). Whether the PMv^DAT^ neurons’ action in the same social behaviors is associated with AVPV/PeN TH neuronal innervation has not been determined.

Our present findings demonstrate that the role of PMv neurons in regulating reproductive physiology is more complex than previously anticipated. While overall PMv^Leprb^ neurons play a significant role in female reproduction, particularly mediating leptin’s permissive effects in puberty (16), PMv^DAT^ cells have been studied primarily in the context of social behavior and aggression (25, 26, 28, 29). The dynamic changes observed in *Slc6a3* gene expression during puberty and in response to nutrition, as well as the developmental difference in the innervation of kisspeptin/TH neurons of AVPV/PeN, suggest that the PMv^DAT^ neurons also play a role in sexual maturation.

## Author contributions

**CSM, CFE** Conceived and designed the experiments. **NB, JDJr, KWW,** generated preliminary data. **CSM, MAS, CNF, TTZ, CMS, LH,** performed the experiments and acquired the data. **CSM, MAS, CB, RF, CFE** analyzed and interpreted the data. **CSM** wrote the manuscript. All authors were involved in revising and approving the manuscript

## Acknowledgements

We thank Dr. Yun-Hee Choi for the design of the DAT riboprobe, and Susan Allen for expert technical assistance. This work was supported by the National Institutes of Health (R01-HD-069702 to CFE, CSM and NB, R21 HD109485 to CFE and CSM), Michigan Nutrition and Obesity Research Center (URM Pilot Grant P30 DK089503 to CSM) CNPq (Brazilian National Council for Scientific and Technological Development fellowship to BCB), the Knut and Alice Wallenberg Foundation (2020.0054) and the Swedish Research Council Distinguished Professorship Grant (2021-00671) to CB, the Coordenação de Aperfeiçoamento de Pessoal de Nível Superior – Brasil (CAPES) – Finance Code 001 (MAS) and by the São Paulo Research Foundation [FAPESP-Brazil, grants number: 13/07908-8 (RF), 15/20198-5 (TTZ) and the National Institutes of Health (R01 DK119169, R56 DK135501, and PO1 DK119130-03 to KWW).

